# Chronic exposure to PFAS triggers systems-level cellular reprogramming independent of their bioaccumulation

**DOI:** 10.1101/2025.07.03.662990

**Authors:** Jenise Z. Paddayuman, Judith R. Cristobal, Luane J. B. Landau, Ashleigh L. Gagnon, Omer Gokcumen, Diana S. Aga, G. Ekin Atilla-Gokcumen

## Abstract

Per- and polyfluoroalkyl substances (PFAS), or “forever chemicals,” are linked to metabolic, immune, and neurotoxic disorders, yet their long-term cellular effects remain unclear. Using a 24-week chronic exposure model with non-transformed human epithelial cells, we examined responses to low, environmentally relevant concentrations of perfluorooctanoic acid (PFOA) and perfluorooctane sulfonic acid (PFOS). Integrated transcriptomic and lipidomic analyses revealed that cellular accumulation was minimal, and molecular changes instead emerged around week 17, marked by activation of oxidative stress responses, cell survival pathways, and lipid metabolism. Our data support a multi-faceted model in which PFAS-induced oxidative stress, mediated by *SESN2* and *SOD3*, alongside increased lipid biosynthesis via SREBP axis and compound-specific disruptions of membrane lipids. These findings highlight the importance of multi-omic, time-resolved approaches in uncovering mechanisms of chronic low-dose chemical exposure and provide a foundation for future *in vivo* studies.

## Introduction

Perfluorooctanoic acid (PFOA) and perfluorooctane sulfonic acid (PFOS) are legacy per- and polyfluoroalkyl substances (PFAS) that are no longer manufactured or used in many countries, yet they extremely persist in the environment (*1, 2*) due to their stability and extensive use (*3*). These compounds also bioaccumulate (*4*) due to widespread exposure (*3, 5, 6*). Because of their remarkable resistance to degradation (*7*), PFOA and PFOS have accumulated in ecosystems over decades, resulting in long-term contamination and detectable levels in both wildlife and human populations. As such, PFAS has been commonly referred to as “forever chemicals” (*7*).

PFOA and PFOS have been detected in various human tissues and are highly bioaccumulative (*8*). Exposure to PFOA and PFOS has been associated with metabolic disorders (*9, 10*), immune dysregulation (*6, 11*), and neurotoxicity (*12–14*). Despite being phased out globally due to health concerns, the molecular mechanisms driving their toxicity remain poorly understood. At the molecular level, PFAS can interact with lipids (*15, 16*), proteins (*17*), and nucleic acids (*18*) as evidenced in biochemical studies. Their amphiphilic structure enables them to interact with lipid membranes, potentially affecting membrane fluidity and permeability (*19, 20*). They can also bind proteins with hydrophobic domains, such as serum albumin (*21*), peroxisome proliferator-activated receptors (PPARs) (*18*), and fatty acid-binding proteins (*22*), potentially interfering with key regulatory functions. Despite these known interactions, the specific cellular pathways and biological processes disrupted by PFAS in living systems remain unclear.

One of the major challenges in understanding the biological pathways affected by PFAS in living systems stems from the discrepancy between laboratory exposure models and real-world environmental exposure. Most experimental studies, including *in vitro* models, employ short-term exposures at relatively high concentrations, often reaching into the high micromolar (10-250 μM) (*23*) concentration range for PFOS and PFOA (5-125 µg/mL), while concentration even at sites of contamination does not exceed ∼150-200 nM for PFOA and PFOS (*2*). This is because little to no cellular response is observed at lower concentrations over short durations of exposure. However, this experimental approach does not accurately reflect the low-dose, chronic exposure scenarios experienced by organisms in the environment. As a result, such models may fail to capture the cumulative or adaptive biological responses that emerge only with prolonged, low-level exposure, limiting the translational relevance of these findings to real-world conditions.

In this study, we employed integrated transcriptomic and lipidomic analyses to investigate the cellular effects of chronic PFOA and PFOS exposure. While previous *in vitro* and *in vivo* studies have used these approaches, they have largely focused on short-term, high-concentration exposures, often reporting conflicting trends in transcriptomic, serum, and cellular lipid profiles. To better reflect real-world exposure, we conducted a time-course experiment using human retinal pigment epithelial cells (hTERT RPE-1), exposing them to environmentally relevant concentrations of PFOA or PFOS over a six-month period. Using RNA sequencing and high-resolution mass spectrometry-based lipidomics, we profiled molecular alterations across time.

Our findings offer new insights into the systems-level response to prolonged, low-concentration PFAS exposure. Rather than a single dominant pathway, PFAS toxicity appears to arise from cumulative and multifactorial disruptions in key cellular processes. This includes dynamic remodeling of the lipidome and sustained activation of oxidative and cellular stress responses. Together, our results provide specific mechanistic hypotheses and a robust framework for future *in vivo* studies aimed at linking genetic and lipid-level responses to PFAS-induced toxicity.

## Results

### A Novel In Vitro Model of Prolonged PFAS Exposure Reveals Limited Cellular Accumulation

To investigate the cellular effects of PFAS at concentrations reflecting realistic environmental exposure, we exposed the human retinal pigment epithelial cell line (hTERT RPE-1) to either 10 nM of PFOA (4.14 µg/L) or 10 nM PFOS (5.00 µg/L), individually, over 24 weeks (**Fig. 1A**). This experimental design provides a suitable framework for modeling the long-term cellular impact of exposure. First, the exposure levels fall within the range detected in contaminated groundwater sources (*2*). Second, hTERT RPE-1 cells are a non-cancerous model that more accurately recapitulates normal cellular homeostasis than commonly used transformed, cancer-derived cell lines, enabling consistent metabolic activity throughout the prolonged exposure period (*15*).

**Fig. 1.**
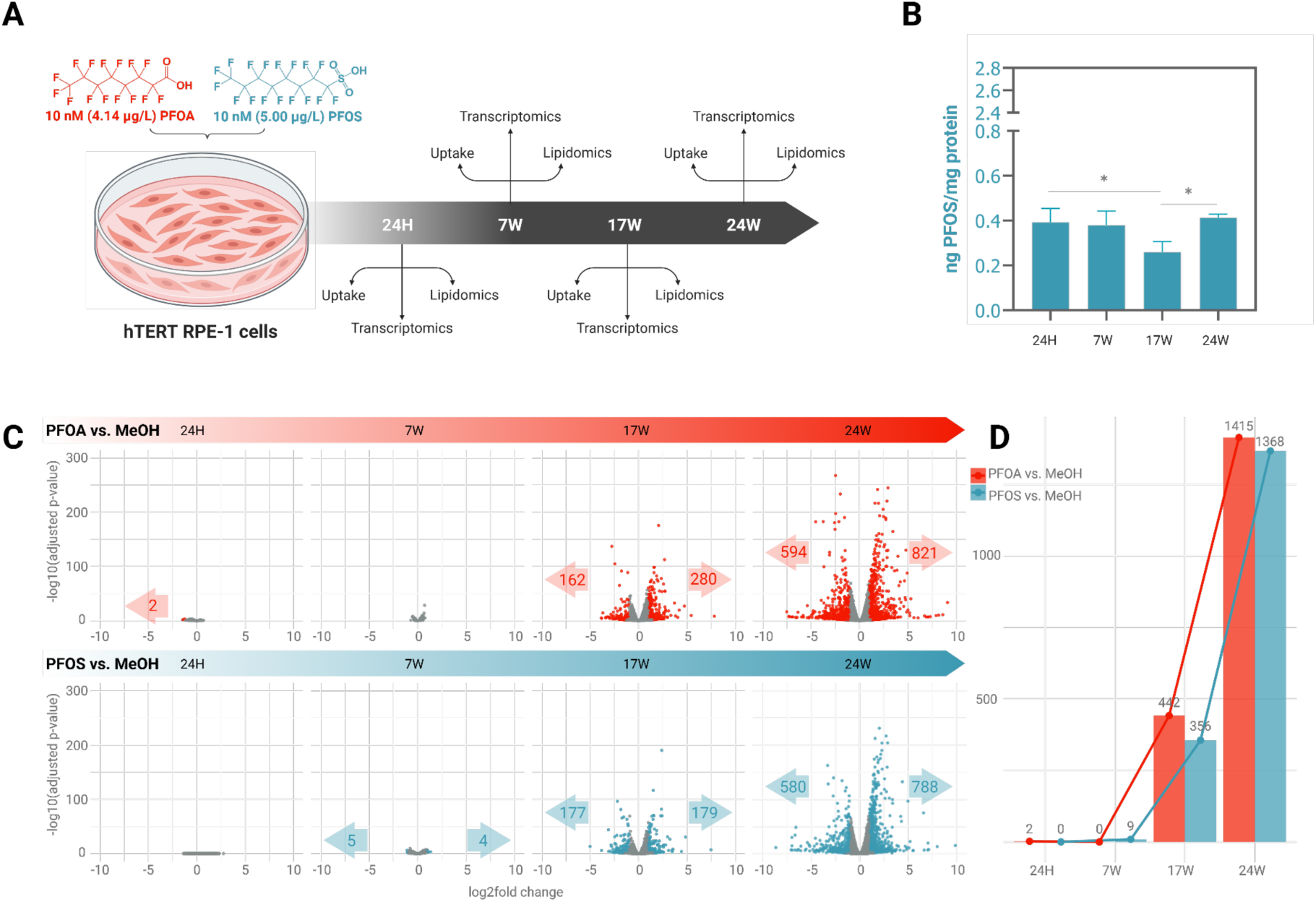
*In vitro* model of chronic PFAS exposure and overview of gene expression changes in hTERT RPE-1 cells. **(A)** Experimental design investigating the uptake, lipidomic, and transcriptomic alterations in human hTERT retinal pigment epithelial (hTERT RPE-1) cells exposed to PFOA (red) and PFOS (blue) over 24 weeks. **(B)** PFOS uptake in hTERT RPE-1 cells. Data are represented as a mean of three independent replicates. P values were calculated using a paired t-test with p<0.05 considered as significant. **(C)** Volcano plots showing differentially upregulated (right arrow) and downregulated (left arrow) genes with prolonged exposure to PFOA and PFOS. Colored dots indicate *p* < 0.05 and |log_2_FC|>1. **(D)** Total differentially regulated genes observed over 24 weeks.

To assess the cellular accumulation of PFAS, we quantified intracellular levels of PFOA and PFOS at 24 hours, 7 weeks, 14 weeks, and 24 weeks (**Methods, Table S1**). Surprisingly, we observed no significant accumulation of either compound over time (**Fig. 1B, Table S2**). Specifically, PFOS exhibited initial uptake within the first 24 hours, reaching a mean concentration of 0.40 ng/mg of protein, followed by a slight decline at 17 weeks and stabilization by 24 weeks (**table S2**). In contrast, we were unable to quantify PFOA as their concentrations at cellular levels were below the limit of detection of our method (LOD = 0.75 ng PFOA/mg protein, **fig. S1 and table S1**). The unexpected observation that PFAS does not accumulate in cells over time suggests that organismal-level bioaccumulation of PFAS may depend on mechanisms beyond direct intracellular uptake, such as accumulation in extracellular components or the presence of multiple cell types that may accumulate PFAS via different mechanisms within complex tissue environments.

### Comparative Gene Expression Profiles Reveal Time-Dependent, System-Level Responses to Prolonged PFAS Exposure

Our findings of minimal or undetectable intracellular bioaccumulation prompted us to investigate whether low-level exposure could still elicit a biological response over time. To address this, we performed comparative transcriptomic analyses across multiple time points under prolonged PFAS exposure (**Fig. 1C, Data S1**). Using a conservative threshold (adjusted *p*-value < 0.05, |log₂FC| > 1; see **Methods**), we observed virtually no differentially expressed genes at the 24-hour and 7-week time points for either PFOA or PFOS. However, at 17 weeks, we detected a sharp increase in transcriptional activity, with 442 and 356 differentially expressed genes for PFOA and PFOS, respectively, compared to methanol-treated controls (**Fig. 1D, Data S1**). This systemic transcriptional response intensified further by 24 weeks, with 1,415 and 1,368 differentially expressed genes for PFOA and PFOS, respectively. These findings suggest that even in the absence of significant intracellular accumulation, prolonged low-level PFAS exposure induces a delayed but robust transcriptomic response, affecting hundreds to thousands of genes over time.

### Cellular Stress is the Major Driver of Convergent Transcriptional Responses to PFAS Exposure

We observed a similar number of differentially expressed genes in response to both PFOA and PFOS treatments. To determine whether these transcriptional changes reflected a shared molecular response, we compared the magnitude and trends in the changes in gene expression upon PFOA and PFOS exposure (**Data S1**). This analysis is critical, as PFOA and PFOS, while structurally related, differ in key biological properties, including half-life (*8*), tissue accumulation, and interaction with host biomolecules (*24*). For example, PFOS exhibits a longer half-life, accumulates more readily in tissues, and binds more strongly to proteins, with well-established effects on lipid homeostasis in liver cells (*8, 25*).

Despite these differences, we found a strong positive correlation in gene expression fold changes between PFOA and PFOS treatments compared to vehicle-treated control cells (Pearson’s r = 0.788; **Fig. 2A**). This high correlation suggests a core transcriptional response to PFOA and PFOS exposure that is largely independent of the specific chemical characteristics of individual compounds.

**Fig. 2.**
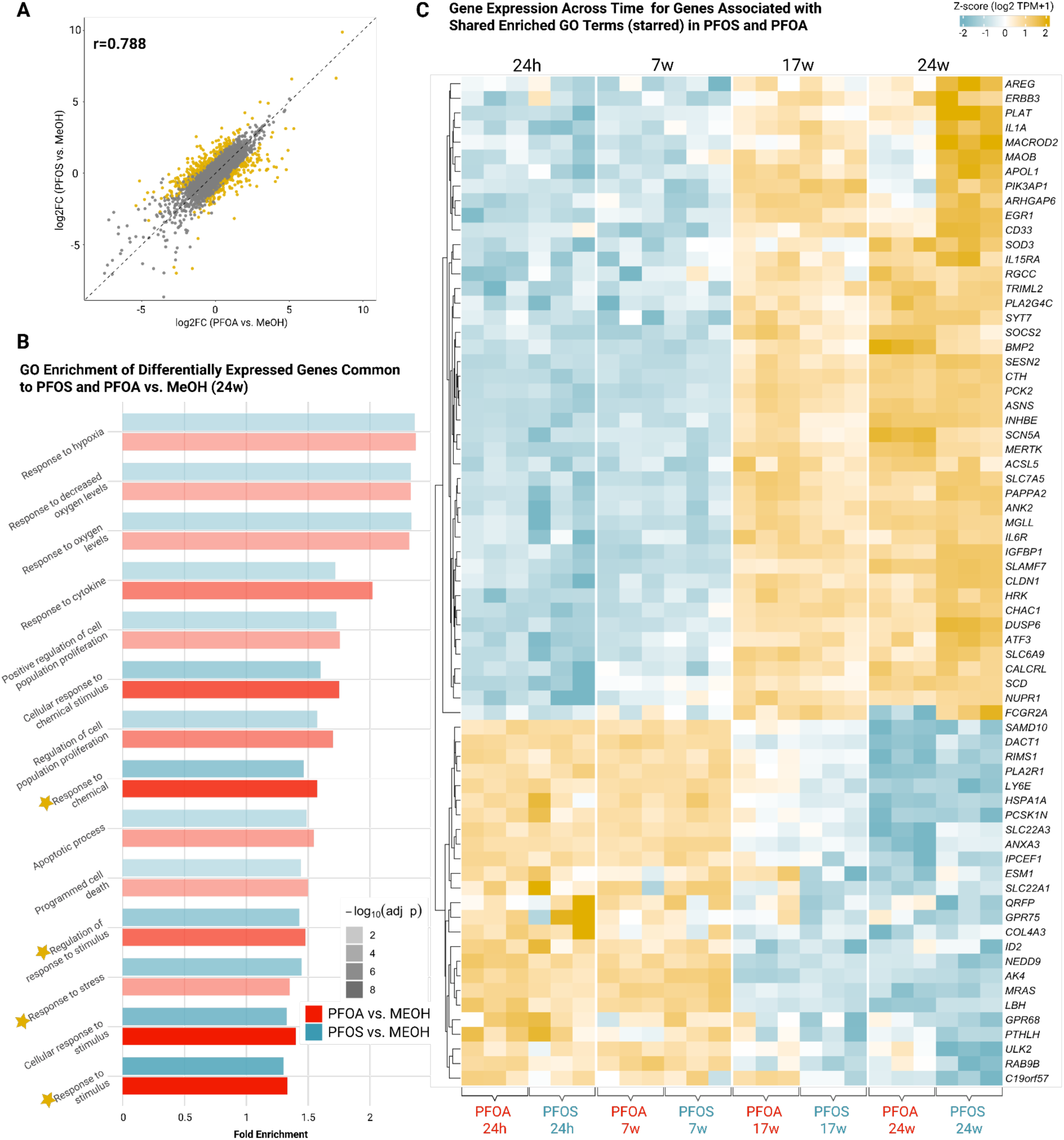
Comparative analysis of gene expression highlighting the transcriptional responses to prolonged PFOA and PFOS exposure of hTERT RPE-1 cells. **(A)** Similarity and consistency in gene expression changes for PFOA and PFOS when compared to the control at 24 weeks. Yellow dots represent genes that are differentially expressed between PFOA and PFOS. **(B)** GO terms related to cellular response and chemical exposure identified from DEGs (*p* < 0.05 and log_2_FC >1) in both comparisons (PFOS vs. MeOH and PFOA vs. MeOH) at 24 weeks. The y-axis shows selected GO terms, the x-axis indicates fold enrichment, and color transparency represents adjusted *p*-values. **(C)** Heat map showing expression trends across time of enriched genes related to the following GO terms: “response to stress”, “response to stimulus”, “regulation of response to stimulus”, and “response to chemical”.

To understand the biological processes underlying this response, we performed Gene Ontology (GO) enrichment analyses (**Methods, Data S2**). The enriched categories shared between PFOA- and PFOS-responsive genes included several biological processes broadly associated with external stress, including oxidative stress, lipid metabolism, and cell death (**Fig. 2B**). To further dissect the pathways involved and to construct a mechanistic framework for the chronic cellular response to PFAS exposure, we conducted a targeted literature review of genes (**Fig. 2C**) contributing to these enriched categories.

Among genes associated with oxidative stress listed in **Fig. 2C**, two stood out as central players: *Superoxide dismutase 3* (*SOD3*) and *Sestrin 2* (*SESN2*), both showing high correlation and consistent upregulation across the exposure period. *SOD3* encodes an extracellular enzyme activated by oxidative stress that converts superoxide radicals, major inducers of cellular damage, into less reactive hydrogen peroxide (*26*). *SESN2*, in contrast, functions intracellularly. It is also induced by oxidative stress and acts as a direct scavenger of reactive oxygen species (ROS), while also regulating broader intracellular stress responses to maintain homeostasis (*27*).

The coordinated upregulation of *SESN2* and *SOD3* exemplifies a robust antioxidant defense response to prolonged PFAS exposure (**Fig. 3, Data S2**). As we expand upon below, oxidative stress likely serves as a central axis that integrates multiple downstream effects of chronic PFAS exposure, including lipid biosynthesis and the activation of cell death pathways.

**Fig. 3.**
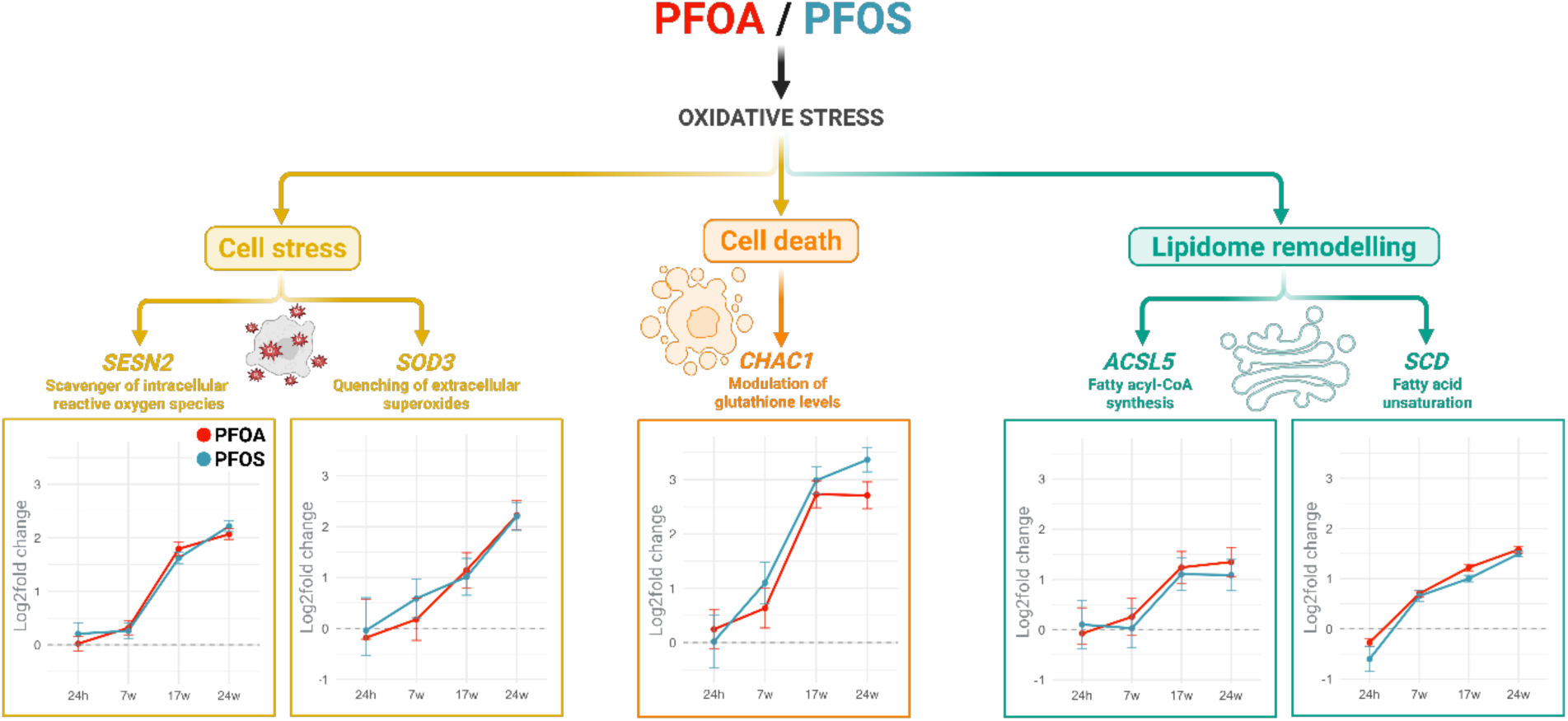
Transcriptomic profiling reveals that chronic exposure to PFOA and PFOS elicits oxidative stress and cellular remodeling responses. Shown are upregulated expression levels (log2fold change) of representative genes involved in oxidative stress [*Sestrin 2 (SESN2), Superoxide dismutase 3 (SOD3)*], cell death [*ChaC glutathione-specific gamma-glutamyl cyclotransferase 1 (CHAC1)*], and lipid metabolism [*Acyl-CoA synthetase long-chain family member 5* (*ACSL5), stearoyl-CoA desaturase (SCD)*]. Data are presented as log2foldchange ± SEM from three independent replicates.

### Lipid and Cell Death-Related Gene Expression Is Driven by PFAS-Induced Oxidative Stress

We next investigated key genes involved in lipid biosynthesis among the enriched categories shared between PFOA- and PFOS-responsive genes (**Fig. 2B and 2C, Data S2**). Two enzymes stood out for their central roles in lipid metabolism: *stearoyl-CoA desaturase* (*SCD*) and *acyl-CoA synthetase long-chain family member 5* (*ACSL5*). *SCD1*, which catalyzes the conversion of saturated fatty acids to monounsaturated fatty acids (*28, 29*), was among the most highly upregulated genes following PFAS exposure (**Fig. 3, Data S2**). This observation is especially compelling in the context of oxidative stress. Prior studies have shown that oxidative stress is associated with the intracellular accumulation of unsaturated fatty acids, a condition that promotes cell death (*30*). Thus, *SCD*-mediated desaturation may act as a protective mechanism to restore lipid homeostasis and reduce lipid-induced apoptotic signaling under oxidative stress conditions.

Similarly, *ACSL5*, which converts long-chain fatty acids (16–20 carbons) into their corresponding acyl-CoA derivatives (*31*), was also upregulated. Long-chain fatty acid oxidation is a well-known source of ROS due to electron leakage during mitochondrial β-oxidation (*32*). By converting free fatty acids into acyl-CoA forms, *ACSL5* may help limit the availability of substrates for ROS-generating oxidation processes, thereby alleviating lipotoxic stress. Together, these changes suggest that shifts in lipid biosynthetic gene expression during chronic, low-level PFAS exposure represent a broader adaptive response to PFAS-induced oxidative stress.

Lastly, we examined *ChaC glutathione-specific gamma-glutamyl cyclotransferase 1* (*CHAC1*), a gene involved in glutathione metabolism and linked to oxidative cell death pathways. *CHAC1* catalyzes the degradation of glutathione into 5-oxo-proline and a cysteinyl-glycine (cys-gly) dipeptide (*33, 34*). Similar to the lipid-related genes, *CHAC1* was significantly upregulated in response to PFAS exposure (**Fig. 2C**, **Fig. 3**). Its increased expression is commonly associated with oxidative stress, where it either reflects or promotes redox imbalance. Thus, *CHAC1* upregulation in our system is consistent with activation of oxidative stress pathways and supports a model in which chronic exposure induces redox disruption. Therefore, upregulation of *CHAC1* contributes to the glutathione depletion, promoting susceptibility to cell death.

It is important to note that, in addition to *CHAC1*, we observed upregulation of a*ctivating transcription factor 3* (*ATF3)*, a transcription factor that is acutely induced under oxidative and cellular stress conditions (**Fig. 2C**) (*35, 36*). Both *CHAC1* and *ATF3* are downstream targets of the integrated stress response pathway and are commonly upregulated in response to ROS accumulation and glutathione depletion (*37*). The coordinated induction of these genes suggests activation of a redox-sensitive transcriptional program, implicating oxidative stress (*33*).

Altogether, gene expressions in **Figure 3** peak at 24 weeks for both PFOA and PFOS treatments, indicating a shared oxidative stress response pattern. Taken together, these findings suggest that cellular stress, lipid metabolism, and cell death pathways are modulated during PFAS exposure as part of an integrated cellular response to oxidative stress.

### PFAS Exposure Rewires the Cellular Lipidome in a Time- and Compound-Specific Manner

Transcriptomic data indicate that oxidative stress-related pathways are broadly impacted by exposure to both PFOA and PFOS, with lipid homeostasis emerging as a central player in this response. Notably, several lipid metabolism-related genes are differentially expressed, prompting us to ask whether these transcriptional changes translate into alterations in the cellular lipidome. To address this, we performed lipidomic profiling following prolonged exposure to PFOA and PFOS.

We targeted members from major mammalian lipid classes (**Fig. 4, fig. S2, and Data S3**). We found that both PFOA and PFOS dramatically reshape the cellular lipidome over time. In contrast to the relatively linear patterns observed in gene expression, lipid abundance followed complex, compound-specific trajectories (**Fig. 4**). Rather than a consistent accumulation or depletion across the lipid family, many lipid species showed dynamic, time-dependent changes, with the 17-week time point emerging as a key inflection point. These effects were especially pronounced in PFOS-exposed cells, where levels of sphingolipids and phosphatidylserines (PS) steadily increased until 17 weeks, then either remained the same or declined to baseline levels by 24 weeks. The dramatic shifts in lipid composition may indicate the presence of multiple, and potentially competing, cellular processes influencing lipid composition in PFAS-exposed cells.

**Fig. 4.**
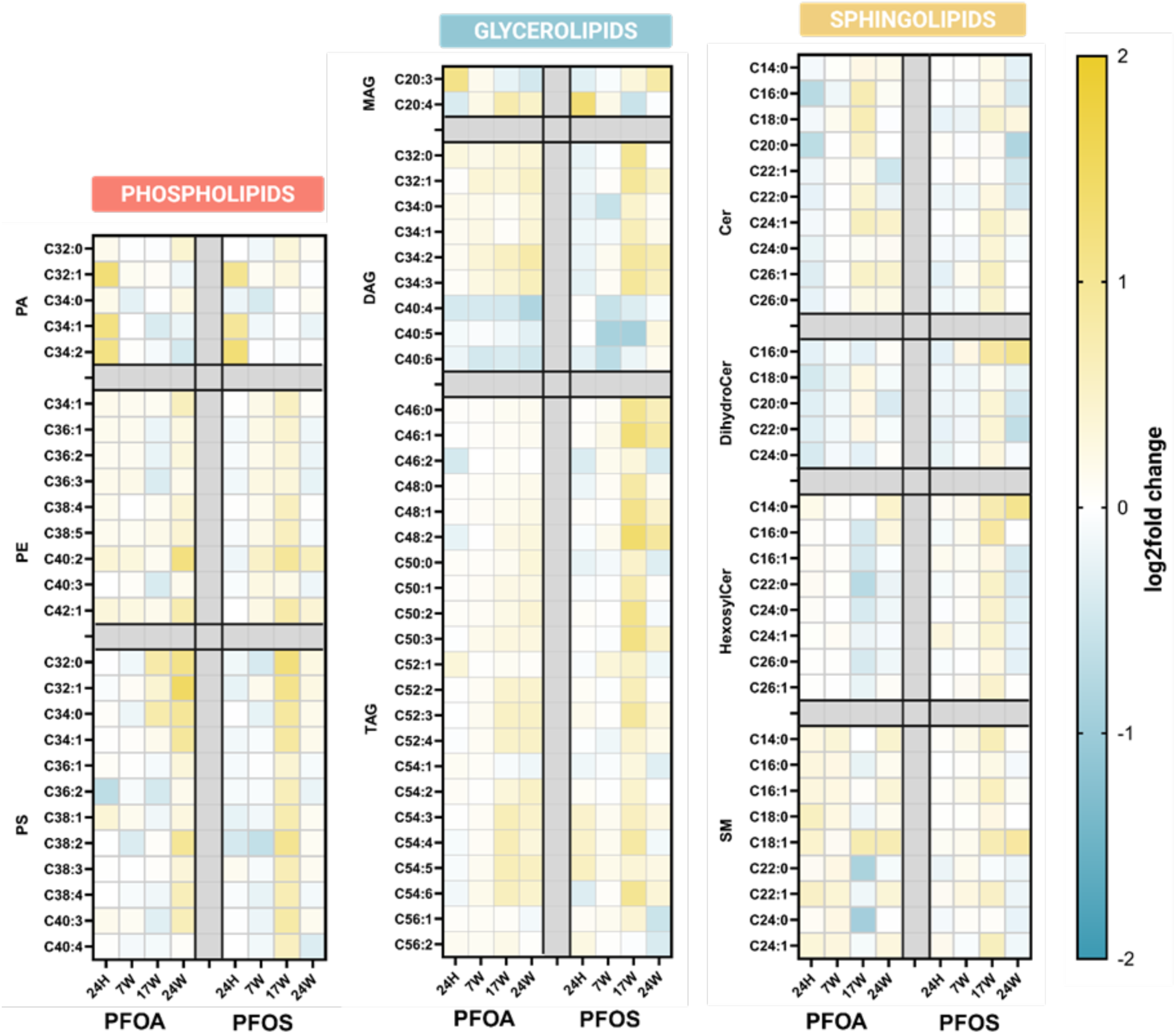
Heatmap representation of lipidomic alterations in hTERT RPE-1 cells following prolonged PFOA and PFOS exposure. The profiles reveal changes across major lipid classes, including phospholipids [phosphatidic acid (PA), phosphatidylethanolamine (PE), and phosphatidylserine (PS)], glycerolipids [monoacylglycerol (MAG), diacylglycerol (DAG), and triacylglycerol (TAG)], and sphingolipids [ceramide (Cer), dihydroceramide (DihydroCer), hexosylceramide (HexosylCer), and sphingomyelin (SM)] in PFOA- and PFOS-treated cells. The log2foldchanges were calculated by normalizing individual abundances to the average of each corresponding time point methanol control. The log2 of this ratio was taken and averaged across three replicates (Refer to **Data S3** for individual values).

### PFAS exposure leads to a general increase in lipid biosynthesis

Previous studies have suggested that PFAS can interact with membranes and interfere with membrane structure and function, triggering compensatory responses through lipid biosynthesis pathways (*38, 39*). One mechanism involves the activation of peroxisome proliferator-activated receptors (PPARs), which regulate transcriptional programs that promote lipid synthesis (**Fig. 5A**) (*39*). Consistent with this, we observed upregulation of multiple PPARs (**Fig. 5B)** and key lipid biosynthesis regulators downstream of PPAR signaling, including *Sterol Regulatory Element Binding Transcription Factor-1 (SREBF1) and Sterol Regulatory Element Binding Transcription Factor-2* (*SREBF2)*.

**Fig. 5.**
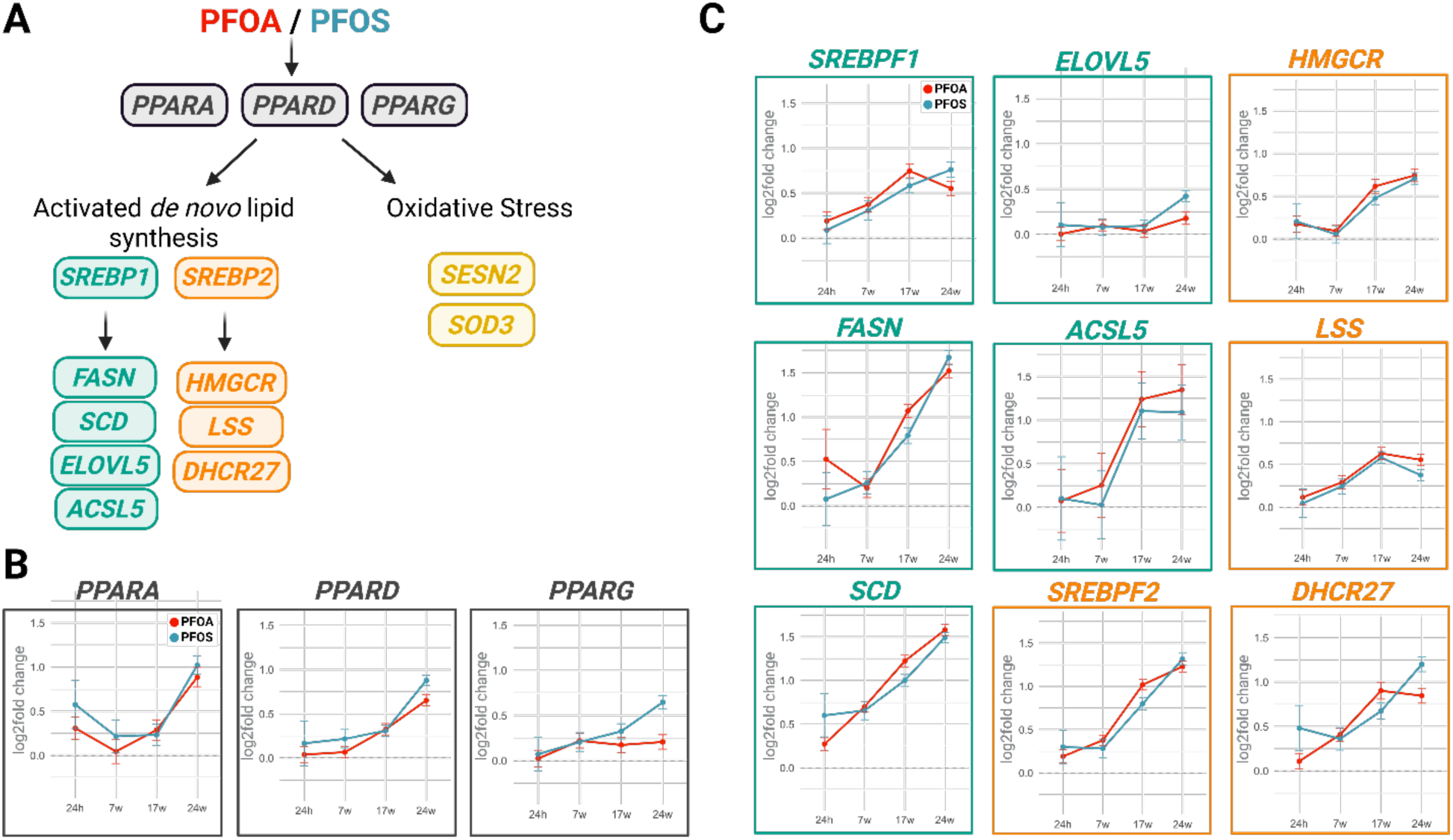
PFOA and PFOS exposure activate PPARs and SREBPs, upregulating key genes in *de novo* lipid biosynthesis. **(A)** Lipid biosynthesis pathways impacted by prolonged PFOA and PFOS exposure lead to oxidative stress, with upregulated *Sestrin 2 (SESN2), Superoxide dismutase 3 (SOD3),* and activated *de novo* lipid synthesis. **(B)** These pathways involve peroxisome proliferator-activated receptors (PPARs) such as PPARA, PPARD, and PPARG. **(C)** The activation of these PPARs leads to transcriptional regulation of genes involved in *de novo* lipid synthesis through *Sterol Regulatory Element Binding Protein-1* (*SREBP1*) [*fatty acid synthase* (*FASN*), *Stearoyl-CoA Desaturase* (*SCD*), *elongation of very long chain fatty acid 5* (ELOVL5), *acyl-CoA synthetase long chain family member 5* (*ACSL5*)], and *Sterol Regulatory Element Binding Protein-2* (*SREBP2*) [*3-hydroxy-3-methyl-glutaryl-coenzyme A reductase* (*HMGCR*), *lanosterol synthase* (*LSS*), and *7-dehydrocholesterol reductase* (*DHCR27*)]. Data are presented as log2foldchange ± SEM from three independent replicates.

The upregulation of *SREBF1* and *SREBF2* are critical in lipid biosynthesis pathways. These genes encode the precursors of the transcription factors *SREBP1* and *SREBP2*, which are master regulators of *de novo* fatty acid, phospholipid, and sterol biosynthesis (**Figs. 5A and 5C, Data S2**) (*40*). Their function depends on spatial translocation from the endoplasmic reticulum membrane to the Golgi apparatus (*41*), where they undergo proteolytic cleavage to generate their active DNA-binding forms. These active forms then translocate to the nucleus to initiate lipid synthesis gene expression. Further, we found a concurrent upregulation of *SREBP Cleavage-Activating Protein* (*SCAP*), which encodes an adaptor protein essential for activation of these two SREBPs, further corroborating the involvement of SREBPs in PFOA- and PFOS-mediated lipid modeling (*41*).

Activation of SREBP downstream targets results in complex alterations in lipid species composition, encompassing a broad range of chain lengths and saturation states, including those involved in glycerolipid and phospholipid synthesis (**Fig. 5A**). Consistent with our expectations, PFAS exposure leads to increased expression of the downstream targets of *SREBP1* (**Fig. 5C**). These targets include key enzymes involved in fatty acid synthesis, activation, elongation, and desaturation. The expression shifts in these enzymes likely underlie the observed increases in glycerolipid and phospholipid species in our lipidomic data. Similarly, *SREBP2* targets were also upregulated, including genes encoding enzymes required for sterol biosynthesis (**Fig. 5C**). Notably, *HMGCR*, which encodes HMG-CoA reductase, the rate-limiting enzyme in sterol synthesis (*42*), was upregulated alongside downstream enzymes involved in sterol ring formation, such as *lanosterol synthase* (*LSS*) (*43*). Together, these findings point to a coordinated increase in lipid production through *de novo* synthesis pathways regulated by the SREBP signaling axis in response to chronic PFOA and PFOS exposure.

### Oxidative stress response contributes to complex lipid composition in PFAS-exposed cells

Despite the global activation of lipid synthesis as a response to PFAS-induced membrane disruption, oxidative stress emerges as a competing force in shaping the lipid landscape of PFAS-exposed cells. As suggested earlier, oxidative stress is a driver of transcriptional changes in response to PFAS exposure, and notably, both membrane disruption and PPAR activation have themselves been shown to exacerbate oxidative stress (*44*). In line with this, the upregulation of oxidative stress response genes such as *SOD3* and *SESN2* may further influence lipid remodeling in exposed cells.

A particularly striking lipidomic signature was the reduction of phosphatidic acid species across both PFOA- and PFOS-treated conditions beyond 24-hour treatment (**Fig. 4**). Phosphatidic acid is a central intermediate in membrane lipid metabolism and a known target of lipid peroxidation under oxidative stress. Its accumulation has been associated with membrane destabilization and increased susceptibility to oxidative damage (*45*). Thus, the observed decrease in phosphatidic acid levels likely represents an adaptive mechanism aimed at mitigating oxidative injury.

In parallel, we observed persistently low levels of long-chain unsaturated diacylglycerols, which serve as precursors for triacylglycerol synthesis (**Fig. 4)**. This trend, combined with the peaking of polyunsaturated triacylglycerols during 17 and 24 weeks (**Fig. 6A**), may reflect another protective cellular strategy, as supported by a previous study (*46*). Specifically, the upregulation of *Diacylglycerol O-Acyltransferase 1* (*DGAT1*) (**Data S2**), which catalyzes triacylglycerol synthesis, could indicate a mechanism to sequester polyunsaturated fatty acid–containing lipids away from membranes. Such sequestration is known to reduce oxidative damage (*47*) during cell death by preventing lipid peroxidation of vulnerable unsaturated fatty acyl chains.

**Fig. 6.**
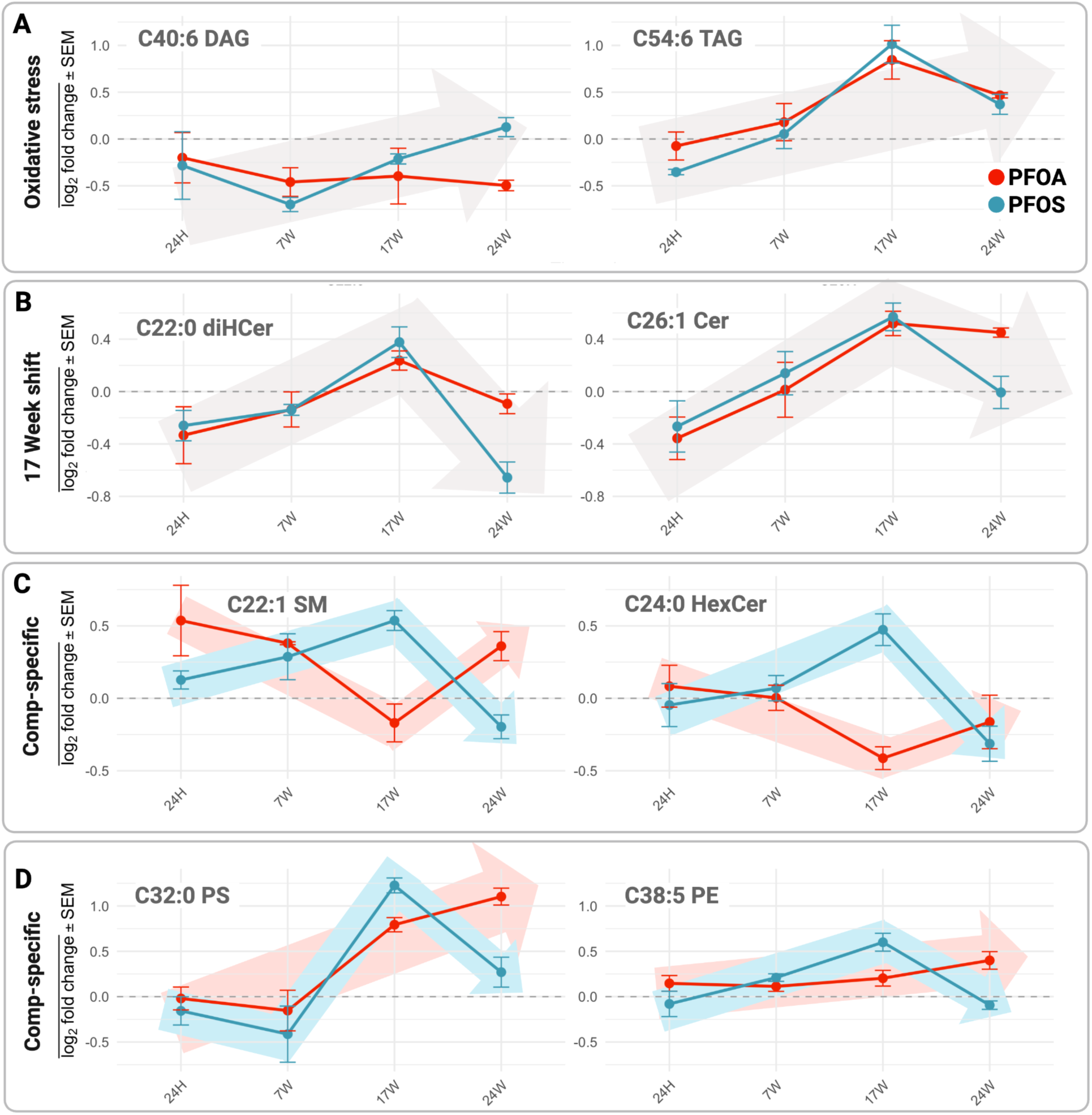
Representative time plots of changes in lipid levels over time following PFOA and PFOS exposure. **(A)** Low levels of long unsaturated diacylglycerols (DAGs) (C40:6) as precursors for triacylglycerol (TAG) biosynthesis, and elevated lipid levels of polyunsaturated TAGs (C54:6) compared to saturated ones, suggest cells are undergoing oxidative stress. **(B)** Observed peaking of lipid levels at 17 weeks for both PFOA and PFOS in dihydroxyceramides (diHCer) (C22:0), and ceramides (Cer) (C26:1). **(C)** Opposite trend on lipid levels observed in PFOA and PFOS in sphingomyelin (SM) (C22:1) and in hexosylceramide (HexCer) (C24:0), **(D)** Compound-specific trend observed in phosphatidylserine (PS) (C32:0), and phosphatidylethanolamine (PE) (C38:5), where PFOA shows upregulation till 24 weeks while PFOS peaks at 17 weeks. Data are presented as log2foldchange ± SEM from three independent replicates.

Taken together, our results suggest that PFOA and PFOS induce a chronic lipotoxic environment characterized by early and persistent neutral-lipid storage and reduction in PAs. This stress-driven remodeling helps explain deviations from the otherwise general trend of increased lipid biosynthesis in PFAS-exposed cells and underscores the complexity of the cellular lipid response under chronic chemical stress.

### Week 17 Marks a Turning Point in Lipid Remodeling and Highlights Distinct Effects of PFOA and PFOS

One of the most notable and unexpected trends we observed was the initial accumulation of multiple lipid species, which peaked at 17 weeks and declined at 24 weeks (**Fig. 6A-B, Data S3**). For example, PFOA exposure led to a sustained buildup of glycerolipids, particularly triacylglycerols, from the early exposure through 24 weeks (**Fig. 4)**. This pattern suggests a diversion of fatty acids into storage lipid droplets that persists for PFOA expression. In contrast, the triacylglycerol levels in PFOS-treated samples declined after 17 weeks, returning to or dropping below baseline levels by 24 weeks. A comparable temporal pattern was observed for ceramides and dihydroceramides, which were elevated at 17 weeks in both PFOS- and PFOA-treated cells but declined by 24 weeks (**Fig. 6B**).

We hypothesized that the exposure-specific, time-dependent differences observed in certain lipid species may also arise from distinct interactions between PFOA or PFOS and cellular membranes. Specifically, the differing chemical properties of their headgroups, carboxylate for PFOA and sulfonate for PFOS, could influence how each compound associates with lipid bilayers, including differences in penetration, orientation, and membrane affinity. These interactions may result in compound-specific patterns of membrane disruption. If this is the case, we would expect to observe differential effects on lipid species that are key structural components of eukaryotic membranes when comparing PFOA and PFOS exposures. Indeed, in PFOS-exposed cells, sphingomyelin and hexosylceramide levels increased over time, peaking at 17 weeks before returning to baseline by week 24. In contrast, PFOA exposure resulted in a marked reduction in both lipid species at 17 weeks, followed by normalization to baseline levels by 24 weeks (**Fig. 6C).** Phosphatidylserines and phosphatidylethanolamines exhibited a distinct temporal pattern. PFOA exposure led to a gradual increase in these lipid species, reaching maximal levels at 24 weeks. In contrast, PFOS exposure induced more dynamic fluctuations, with levels peaking at 17 weeks before declining **(Fig. 6D**).

Overall, our findings suggest multiple novel hypotheses concerning the compound-specific effects of PFAS on membrane structure and lipid homeostasis. These insights lay the groundwork for future functional studies aimed at dissecting how distinct PFAS compounds interact with cellular membranes and how the timing of cellular responses influences long-term exposure.

## Discussion

Our study reveals that prolonged exposure to environmentally relevant concentrations of PFOA and PFOS elicits robust, time-dependent cellular responses in the absence of significant intracellular PFAS accumulation. Using an *in vitro* model with non-transformed epithelial cells, we demonstrated that molecular responses to low-dose PFAS exposures unfold gradually, with delayed but coordinated alterations in gene expression and lipid composition emerging primarily after 17 weeks of exposure. These responses converge on oxidative stress, lipid metabolism, and membrane remodeling.

We show that both PFOA and PFOS activate PPAR signaling pathways and induce oxidative stress, as evidenced by the upregulation of canonical antioxidant genes such as *SESN2* and *SOD3*. This stress response, in turn, modulates transcriptional reprogramming of lipid biosynthetic pathways, marked by the activation of *SREBP1/2* targets and increased expression of key enzymes involved in lipid synthesis. However, lipidomics reveals that this transcriptional drive toward lipid production is counterbalanced by stress-adaptive lipid remodeling, including the sequestration of polyunsaturated fatty acids into neutral lipids and the depletion of oxidation-prone lipid intermediates such as phosphatidic acid.

Importantly, we uncovered a critical temporal inflection point at 17 weeks, where lipid profiles undergo substantial shifts and begin to diverge between PFOA- and PFOS-treated cells. These compound-specific trends, particularly in sphingolipids and membrane phospholipids, suggest that subtle differences in PFAS structure and properties can lead to distinct biophysical interactions with membranes, producing divergent cellular outcomes over time.

Altogether, our findings propose a model in which chronic PFAS exposure triggers a multilayered, systems-level response involving oxidative stress, transcriptional regulation of lipid metabolism, and membrane remodeling. These results provide a molecular framework for understanding how chronic, low-level chemical exposures can disrupt cellular homeostasis and shape long-term physiological outcomes. Future work investigating how distinct PFAS interact with membrane components *in vivo* and across cell types will be critical for elucidating their long-term health impacts.

## Materials and Methods

### Materials

hTERT RPE-1 cells were purchased from the American Type Culture Collection (Manassas, VA). Dulbecco’s modified Eagle media (DMEM)/Ham’s F-12 50:50, fetal bovine serum (FBS), trypsin, and penicillin/streptomycin (p/s) antibiotic cocktail were purchased from Corning (Corning, NY). Micro bicinchoninic acid (BCA) assay kit was purchased from G Biosciences (St. Louis, MO).

PFOA and PFOS were purchased from Millipore Sigma (Burlington, MA). ^13^C-labelled standards, including M8PFOA, M8PFOS, M4PFOA, and M4PFOS, were purchased from Wellington Laboratories (Guelph, ON). Ammonium acetate was purchased from J.T. Baker (Randnor Township, PA). Liquid chromatography-mass spectrometry (LC-MS) grade methanol (MeOH), isopropanol, and acetonitrile were purchased from Millipore Sigma (Burlington, MA). LC-MS grade chloroform was purchased from Honeywell Research Chemicals (Morris Plains, NJ). Type I water (18.2 (18.2 MΩ cm) was generated using a Barnstead Nanopure Diamond (Waltham, MA) purification system. LC-MS column and guard column were obtained from Phenomenex (Torrance, CA).

MassHunter^TM^ Qualitative Analysis, which was utilized for data analysis, was obtained from Agilent Technologies (Santa Clara, CA).

### Methods

#### Cell culture, PFAS exposure, and cell collection

The culturing, PFAS exposure, and subsequent collection of hTERT RPE-1 cells were conducted following a recently published protocol (*15*). Briefly, hTERT RPE-1 cells were cultured in Dulbecco’s Modified Eagle Medium/Nutrient Mixture F-12 (DMEM/F-12) with L-glutamine and 15 mM HEPES. The media was supplemented with 10% (v/v) fetal bovine serum (FBS) and 1% (v/v) penicillin/streptomycin (p/s). Cells were grown at 37°C and 5% CO2 until they reached 80-90% confluency for use. Cells were treated with methanol (vehicle control), 10 nM PFOA, or 10 nM PFOS in separate 10-cm dishes and sub-cultured biweekly for 24 weeks. At designated time points (24 hours, 7 weeks, 17 weeks, and 24 weeks), cells at 80–90% confluency were collected. Adherent cells were collected and centrifuged at 300 rcf for 5 mins. at 4°C. The media was decanted, and the cells were washed three times with cold 1x phosphate-buffered saline (PBS) to remove residual media using the centrifugation protocol previously described. The cell pellet was resuspended in 500 mL of cold 1x PBS. For PFAS uptake analysis, 50 μL aliquots were set aside for protein quantification. Cell pellets were stored at −80°C for subsequent analysis: bioaccumulation, lipidomics, and transcriptomics. At every exposure time point, each treatment group included at least triplicate samples (n=3) for every analysis.

#### PFAS Extraction and Liquid Chromatography-Tandem Mass Spectrometry (LC-MS/MS) Data Acquisition for PFAS Uptake

The PFAS-treated cells were extracted and analyzed following a published protocol (*12*). Cell pellets treated with PFOS and PFOA, along with the vehicle control, were resuspended in 1 mL of cold methanol spiked with corresponding mass-labelled PFAS standards (M8PFOS and M8PFOA) as surrogates. The cell suspension was sonicated using a probe sonicator set at 40% power for 30 seconds while kept on a cold metal block. After sonication, samples were centrifuged at 16,900 rcf for 12 minutes at 4°C. Carefully, 900 μL of the resulting supernatant was transferred into a 1 mL dram vial. The remaining pellets were subjected to a second extraction by adding 1 mL of methanol and repeating the sonication and centrifugation steps. The supernatants from both extractions were combined and evaporated to dryness using a nitrogen evaporator. Final extracts were reconstituted with the mobile phase system of 50:50 (A) 5mM ammonium acetate in water with 5% acetonitrile, pH 3.8; and (B) 50:50 methanol: acetonitrile (v/v) spiked with 50 ppb of internal standards, M4PFOS and M4PFOA. PFOA and PFOS samples were analyzed in negative mode as [M-H]^-^ or [M-COOH]^-^ adducts via liquid chromatography-tandem mass spectrometry (LC-MS/MS) using a Thermo TSQ Quantum Ultra^TM^ triple quadrupole mass spectrometer and Agilent 1200 liquid chromatograph. Separation of each 10 μL injection was achieved using a Raptor C18 analytical column (100 mm, 2.7 μm) (Restek Corp., Bellefonte, PA, USA), equipped with a 5 mm guard column containing identical stationary phase material, using an elution gradient and 0.37 mL/min flow rate. The gradient profile consisted of 50% (B) ramped to 95% (B) over 8 min and was held for 3 mins before returning to the starting mobile phase of 50% (B). The total run time for each injection was 18 min. The mass spectrometer was operated in selected reaction monitoring mode, and two transitions were monitored for each analyte to ensure specificity.

#### Limit of Quantification (LOQ) and Limit of Detection (LOD) Determination for PFOS and PFOA

Ten 10-cm plates of confluent hTERT RPE-1 cells were collected, protein quantified, and extracted following the above protocol. To create the matrix-matched calibration curves, for each PFOA and PFOS, seven-point calibration with replicates (n=3) were prepared using serial dilution starting with 1 ppb to 64 ppb in diluent matrix containing 50 ppb M8PFOA and M8PFOS as an internal standard for normalization. The slope (S) was then obtained from the calibration curves (**fig. S1**). For the lowest concentration (1 ppb), seven replicates were made to get the standard deviation of the response (Sy). Prepared solutions were analyzed using LC-MS/MS as described above. Finally, LODs and LOQs were calculated using the formula LOD = 3(Sy/S) and LOQ = 10(Sy/S) (**table S1**).

#### RNA extraction, library preparation, sequencing, and analysis

Frozen cell pellets consist of MeOH control, 24 hours, 7 weeks, 17 weeks, and 24 weeks (n=5) that were previously stored at -80 °C and were transported on dry ice to Azenta Life Sciences for RNA extraction and RNA-seq analysis. Azenta’s RNA-seq workflow consists of five steps, including experimental design, extraction, library preparation, sequencing, and data analysis. RNA from cells was extracted, and RNA molecules of interest were purified, ensuring high quality and sufficient quantity prior to proceeding with analysis. After which, single-stranded RNA was converted to double-stranded complementary DNA (cDNA) strands in a reverse transcription reaction. Sequencing adapters and barcodes were then added to create RNA-seq libraries that were subsequently analyzed with Next Generation Sequencing (NGS) using Illumina NovaSeq^TM^6000 or Illumina Hiseq^®^X. Data was then evaluated, and biologically relevant information was extracted. Briefly, sequencing reads were pre-processed and trimmed to remove adapter sequences and low-quality bases. Trimmed reads were then mapped to the reference genome, followed by expression quantification and TPM calculation. Differential expression analysis was performed using DESeq2 (*48*), comparing treatments to control in the following contrasts: PFOA vs. MeOH, PFOS vs. MeOH, and PFOA vs. PFOS. Note that MeOH 24H was used as the control for all time point comparisons involving both PFOA and PFOS. GO enrichment analysis was conducted for significant genes (|log2foldchange|>1 & *p* < 0.05) in each comparison using gProfiler (*49*). To investigate shared gene expression patterns between PFOA and PFOS at 24 weeks, we identified DEGs common to both PFOS vs. MeOH and PFOA vs. MeOH comparisons at 24w and performed GO enrichment analysis on this shared gene set.

#### Lipid Extraction and Liquid Chromatography-Mass Spectrometry (LC-MS) Data Acquisition for Lipidomics Analysis

Lipid extraction, sample preparation, and analysis were carried out as described in a published protocol (*50*). Cell pellets were resuspended in 550 µL of cold 1x PBS, and 50 µL of aliquots were set aside for protein quantification. An additional 500 mL of cold 1x PBS was added, and the cell pellets were transferred to a 7 mL Dounce homogenizer. To the suspension, 1 mL of cold methanol and 2 mL of cold chloroform were added, and the mixture was homogenized with 30 strokes. The homogenized solution was transferred to a 2-dram vial and centrifuged at 500×g for 10 mins at 4°C. The chloroform layer, containing the lipids, was carefully extracted and transferred to a new dram vial. This layer was evaporated to dryness using a nitrogen evaporator and subsequently stored at −80°C until further analysis by liquid chromatography-mass spectrometry (LC-MS).

As internal standards, lipid extracts were spiked with C17:0 Ceramide (C17-Cer), C17 Sphingomyelin, d^9^-oleic acid, C17 Glucosyl(b) Ceramide (C17 GluCer), C39:0 triacylglycerol (C39-TAG), C18-d^7^-sphingosine as internal standards and reconstituted with chloroform. Analysis was carried out in negative and positive mode via Agilent 1260 HPLC in tandem with an Agilent 6530 Jet Stream ESI-QToF-MS system. Separation of each 15 μL injection was achieved using Gemini C18 column (5 μM, 4.6 mm × 50 mm) for negative mode, while Luna C5 column (5 μM, 4.6 mm × 50 mm) for positive mode. The mobile phase system consisted of: (A) 95:5 water: methanol and (B) 60:35:5 isopropanol:methanol:water for both modes, with the addition of ammonium hydroxide for negative, while the addition of formic acid and ammonium formate for positive. The gradient profile consisted of 0% (B) ramped to 100% (B) over 65 mins. and was held for 7 mins before returning to the starting mobile phase of 0% (B). The total run time for each injection was 80 mins. Lipid species were targeted by extracting the corresponding *m/z* for each ion in MassHunter^TM^ Qualitative Analysis software (v11). Peak areas for each ion were integrated and represented as abundance. Relative abundances used in heatmaps for the PFAS treatments (n=3) were normalized to the corresponding time point control.

### Statistical Analysis

All statistical analyses were performed using the GraphPad Prism (v10.4.2) or R software (v4.2.1). Data presented are an average of three independent replicates. P values were calculated using a paired t-test with p<0.05 considered as significant. In GO enrichment analysis, |log2foldchange|>1 & p < 0.05 were set as threshold for significant genes gProfiler.

## Acknowledgments

The authors would like to acknowledge UB RENEW Institute for the use of the analytical instrumentation and staff support.

## Author contributions

Conceptualization: JP, JRC, OG, DSA, GEAG

Methodology: HP, FTGS, CW, JRK

Investigation: JP, JRC, LBL, AG

Visualization: JP, JRC, LBL

Supervision: OG, DSA, GEAG

Writing—original draft: JP, JRC, OG, DSA, GEAG

Writing—review & editing: JP, JRC, LBL, OG, DSA, GEAG

## Data and materials availability

All data are available in the main text or the supplementary materials.

## Supplementary Materials

**Table S1.** Matrix limit of detections (LODs) and limit of quantifications (LOQs) for hTERT RPE-1. LODs and LOQs were calculated using the formula LOD = 3(Sy/S) and LOQ = 10(Sy/S) and reported in ppb and ng/mg protein units. The standard deviation of the response (Sy) was obtained from the lowest concentration (n=7) while the slope (S) was obtained from the calibration curve shown in Figure S1.

| PFAS | LOD |  | LOQ |  |
| --- | --- | --- | --- | --- |
|  | ppb | ng/mg protein | ppb | ng/mg protein |
| PFOA | 0.8634 | 0.7508 | 2.8780 | 2.5026 |
| PFOS | 0.0709 | 0.0616 | 0.2362 | 0.2054 |

**Table S2.** Average PFOA and PFOS uptake results +/- standard deviation in hTERT RPE-1 cells. Data are obtained from three independent replicates.

| Time Points | PFOA, ng/mg protein | PFOS, ng/mg protein |
| --- | --- | --- |
| 24 hours | < LOD | 0.39 ± 0.06 |
| 7 weeks | < LOD | 0.38 ± 0.06 |
| 17 weeks | < LOD | 0.26 ± 0.05 |
| 24 weeks | < LOD | 0.42 ± 0.01 |

**Figure S1.**
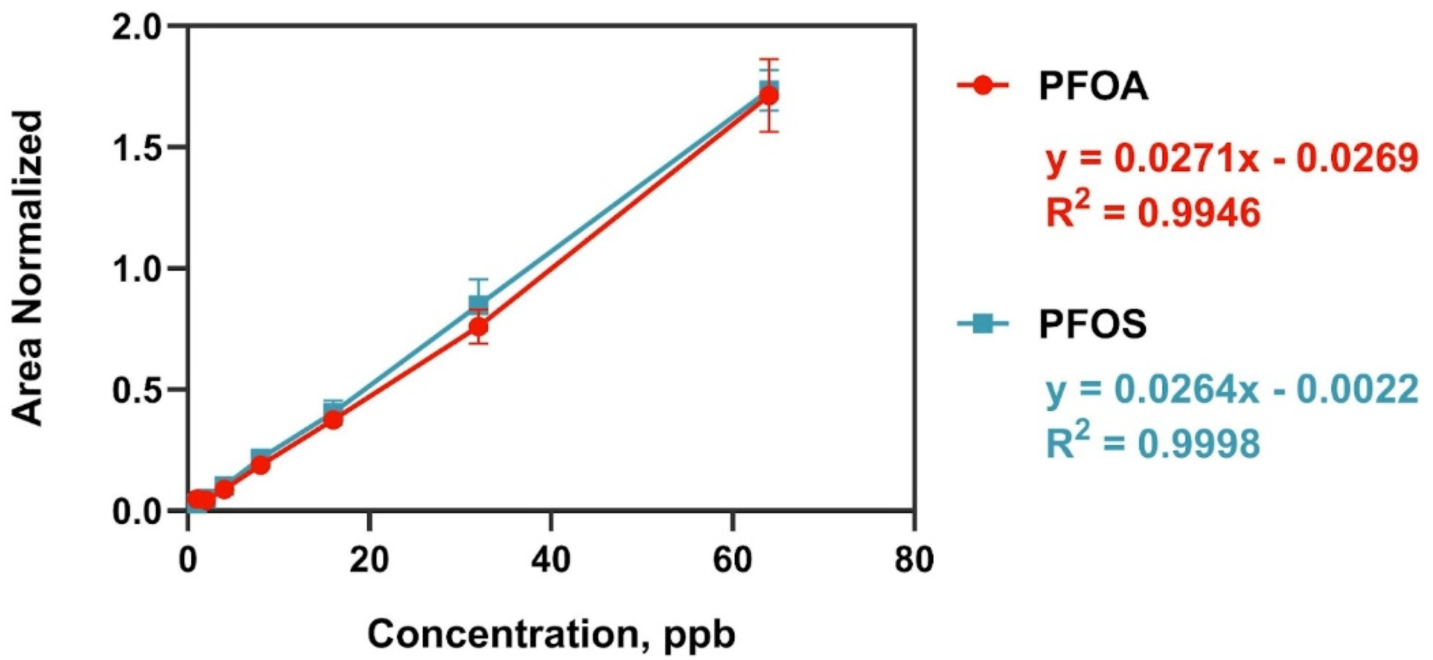
Matrix-matched calibration curves for PFOA and PFOS in hTERT RPE-1 cells. Seven-point calibration with replicates (n=3) were independently prepared for PFOA and PFOS using serial dilution of the highest concentration solution to cover 1 ppb to 64 ppb standards in matrix diluent containing 50 ppb of the corresponding isotopically labeled internal standard (M8PFOA for PFOA and M8PFOS for PFOS) for normalization.

**Figure S2.**
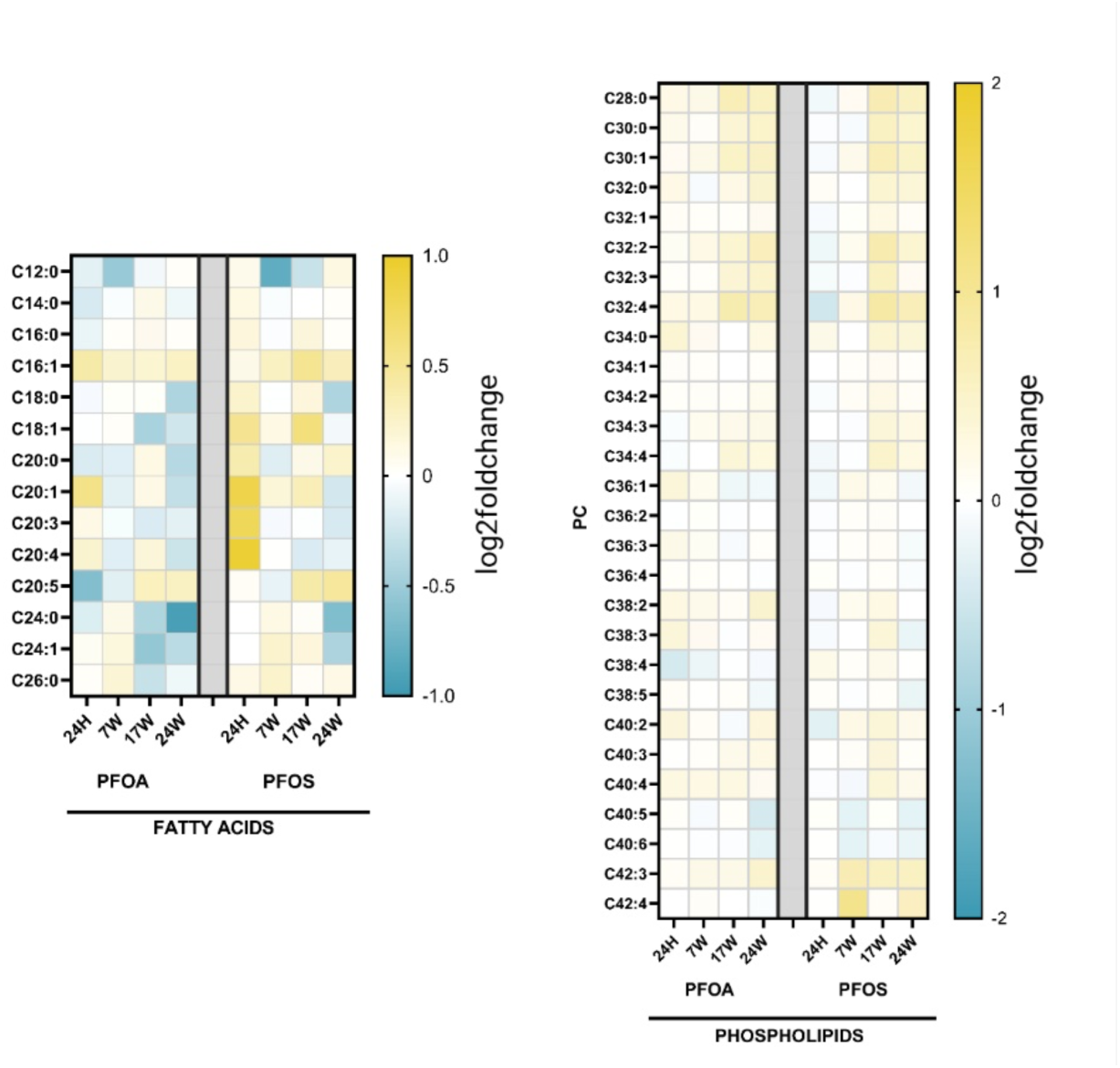
Heatmap of fatty acid and phosphatidylcholines (PC) alterations in hTERT RPE-1 cells after prolonged PFOA and PFOS exposure. Data are shown as log2foldchanges with respect to corresponding control group per time point.

**Data S1. (separated file)**

Comprehensive summary table containing all differential expression results from transcriptomic analyses comparing PFOA and PFOS exposures to MeOH controls across multiple time points. Provided as a separate Excel file.

Excel File: Merged_DESeq2_Data

**Data S2. (separated file)**

Gene ontology (GO) enrichment results for differentially expressed genes (|log₂FoldChange| > 1, *p* < 0.05) from both PFOA vs. MeOH and PFOS vs. MeOH comparisons at the 24-week time point. Results are merged into a separate excel file.

Excel File: 24w_GO_terms_MERGED_PFOS_vs_MeOH &_PFOA_vs_MeOH.csv

**Data S3. (separated file)**

Targeted lipidomics of PFOA and PFOS-treated cells. Relative abundance was calculated as the ratio of each lipid’s abundance to the mean abundance of the control group at the corresponding time point. The log₂ of this ratio was then used to compute the log₂ fold change. This is provided as a separate excel sheet.

Excel File: Targeted_Lipidomics

